# PerMemDB: a database for eukaryotic peripheral membrane proteins

**DOI:** 10.1101/531541

**Authors:** Katerina C. Nastou, Georgios N. Tsaousis, Vassiliki A. Iconomidou

## Abstract

The majority of all proteins in cells interact with membranes either permanently or temporarily. Peripheral membrane proteins form transient complexes with membrane proteins and/or lipids, via non-covalent interactions and are of outmost importance, due to numerous cellular functions in which they participate. In an effort to collect data regarding this heterogeneous group of proteins we designed and constructed a database, called PerMemDB. PerMemDB is currently the most complete and comprehensive repository of data for eukaryotic peripheral membrane proteins deposited in UniProt or predicted with the use of MBPpred – a computational method that specializes in the detection of proteins that interact non-covalently with membrane lipids, via membrane binding domains. The first version of the database contains 231770 peripheral membrane proteins from 1009 organisms. All entries have cross-references to other databases, literature references and annotation regarding their interactions with other proteins. Moreover, additional sequence annotation of the characteristic domains that allow these proteins to interact with membranes is available, due to the application of MBPpred. Through the web interface of PerMemDB, users can browse the contents of the database, submit advanced text searches and BLAST queries against the protein sequences deposited in PerMemDB. We expect this repository to serve as a source of information that will allow the scientific community to gain a deeper understanding of the evolution and function of peripheral membrane proteins via the enhancement of proteome-wide analyses.

The database is available at: http://bioinformatics.biol.uoa.gr/db=permemdb

## 1. INTRODUCTION

One universal feature of all cells, upon which their structure and majority of functions rely, are membranes [1]. Membranes serve as permeable barriers for the entire cell or certain subcellular structures and are associated with the majority of cellular proteins [2]. Signal transduction, molecular and ion transport, enzymatic activity and cell adhesion are among the most important functions performed by membrane proteins [3, 4]. Membrane proteins can be classified into two broad categories based on the nature of membrane-protein interactions; integral and peripheral membrane proteins [1, 5].

Peripheral membrane proteins interact indirectly with membrane proteins or lipids via non-covalent interactions [6-8] and are critical due to the numerous cellular functions in which they participate [9, 10]. Peripheral proteins also possess domains that permit the specific or non-specific interaction with membrane lipids, to perform functions related to signal transduction and membrane trafficking [11, 12]. These domains allow the identification and classification of these proteins [13] and have been exploited for the development of three computational methods for the detection of peripheral membrane proteins in proteomes [14-16]. Among these three methods, MBPpred has the most extended library of pHMMs and detects proteins that possess 18 domains with experimentally validated interactions with membrane lipids [16].

To this day, two databases have been developed that contain data for specific subgroups of peripheral membrane proteins. The first one is OPM [17], a database dedicated to the optimization of the spatial arrangement of protein structures in lipid bilayers and contains data for a substantial number of peripheral proteins derived from PDB [18]. The other one, MeTaDoR [19], is a decade-old online resource devoted specifically to peripheral proteins with membrane targeting domains and includes structural and sequence data. However, OPM contains only a collection of structural data on peripheral membrane proteins and MetaDoR’s online interface has ceased functioning since 2014. At present there is no special-purpose biological database for peripheral membrane proteins available. This fact urges the need for a thorough data collection and more rigorous studies regarding this protein group.

In this study we addressed this need through the construction of PerMemDB, a database for peripheral membrane proteins in eukaryotes. This repository currently holds data on peripheral membrane proteins, deposited in UniProt [20] or predicted with the use of MBPpred [16] for all eukaryotic reference proteomes.

## 2. METHODS

The development of PerMemDB, was based on three data collection approaches. Firstly, the available proteins from UniProt [20] with subcellular location “Peripheral membrane protein [SL-9903]” were isolated for all eukaryotic reference proteomes via programmatic access to the UniProt API [21]. Then, peripheral proteins with Membrane Binding Domains (MBDs) that interact directly with membrane lipids were retrieved, using MBPpred [16], a prediction method developed in our lab, that identifies these proteins from their sequence via a library of profile HMMs, specific for 18 MBDs. For this purpose, FASTA formatted files [22] for all eukaryotic reference proteomes were downloaded automatically from UniProt and used as input files for the stand-alone local version of MBPpred. Finally, in order to collect peripheral membrane proteins that interact indirectly with the membrane, all non-transmembrane interactors of transmembrane proteins (subcellular location: “Multi-pass membrane protein [SL-9909]” or “Single-pass membrane protein [SL-9904]” or “Single-pass type I membrane protein [SL-9905]” or “Single-pass type II membrane protein [SL-9906]” or “Single-pass type III membrane protein [SL-9907]” or “Single-pass type IV membrane protein [SL-9908]”) for eukaryotic reference proteomes were collected programmatically and annotated, also, as peripheral. Figure 1 shows the pipeline for the retrieval of all protein datasets. Seven final subsets of unique peripheral proteins from these three sources were created and stored in PerMemDB (Table 1).

**Table 1.** The subsets of peripheral membrane proteins stored in PerMemDB, grouped by the source(s) from which they were isolated. A description of each subset is given in the last column.

| Set | Name | Description |
| --- | --- | --- |
| 1 | UniProt_only | Entries found only in UniProt, with SL-9903 |
| 2 | MBPpred_only | Peripheral membrane proteins retrieved only with the use of MBPpred |
| 3 | TMint_only | Only non-transmembrane interactors of transmembrane (TM) proteins |
| 4 | UniProt_MBPpred | Entries with SL-9903 which were also retrieved with the use of MBPpred but were not found to interact with TM proteins |
| 5 | UniProt_TMint | Entries with SL-9903 which were also interactors of TM proteins, but were not detected with MBPpred |
| 6 | MBPpred_TMint | Peripheral membrane proteins retrieved with the use of MBPpred, which were also interactors of TM proteins, but were not reported as peripheral in UniProt |
| 7 | UniProt_MBPpred_TMint | Entries with SL-9903, retrieved with MBPpred and found to interact with TM proteins |

**Figure 1.**
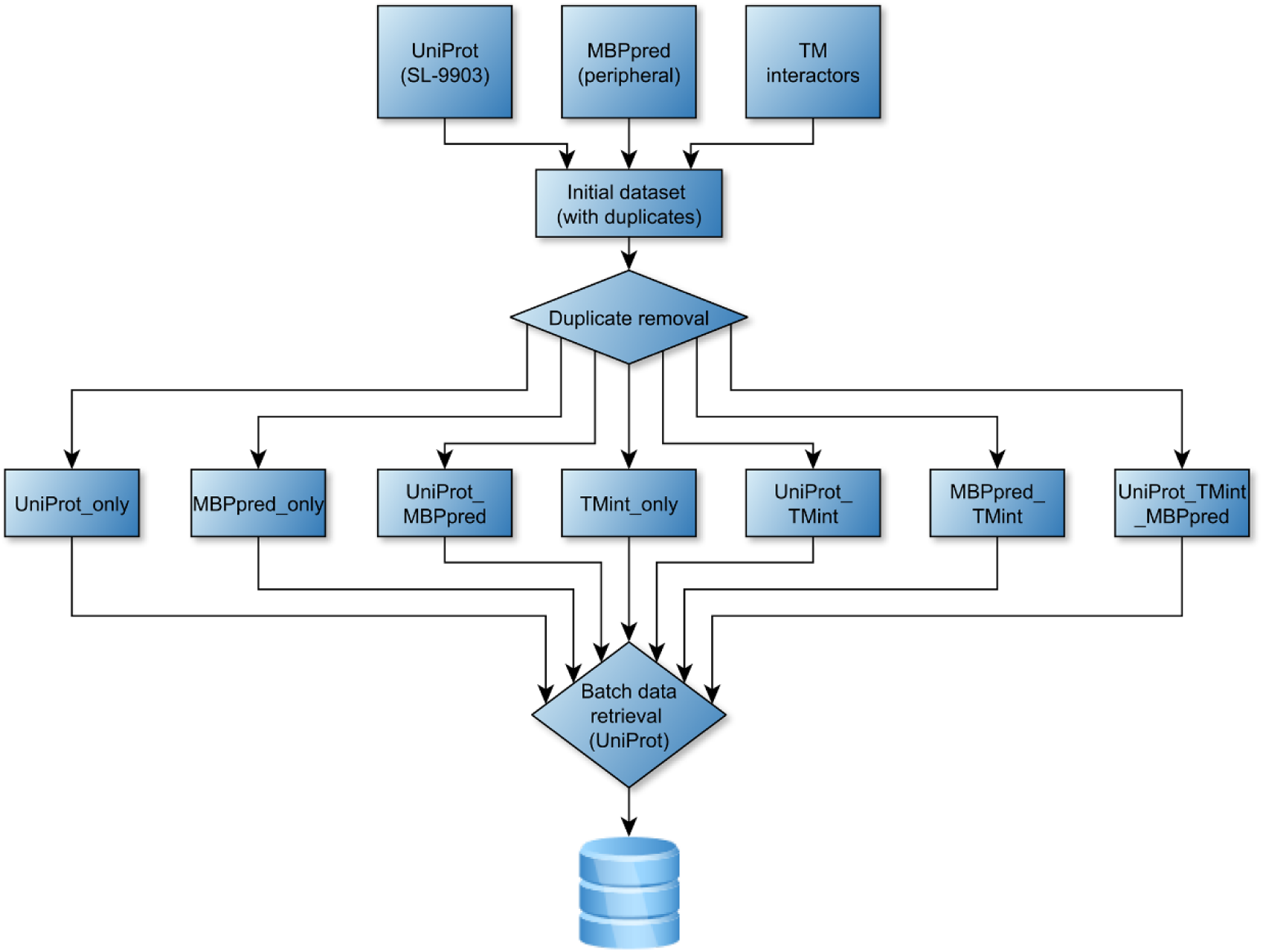
The data retrieval pipeline. Proteins with subcellular location “Peripheral membrane protein” were retrieved from UniProt, “peripheral proteins” with MBDs were identified with the use of MBPpred and non-transmembrane interaction partners of transmembrane (TM) proteins were isolated from UniProt. Scripts were written in Perl and Python for the automated retrieval of all entries with the required information and of all fasta files that were used as input for MBPpred. An initial dataset of protein entries was created after the merge of all aforementioned lists. After a comparison of the three lists, seven non-overlapping datasets were created, that would provide the final set of proteins to be stored in the database. The contents of each dataset are described in Table 1. Python scripts were written to recover data from UniProt for all protein entries and were further manipulated in order to be stored in a relational database.

A web application for PerMemDB has been developed. A mySQL database system was used to store all protein data in a relational database and serves as the first layer of the application. The second layer is a Node.js application server that receives user queries to the database and returns data to the web browser. The web interface is based on modern technologies (HTML5, CSS3 and Javascript) and can be viewed from any screen size (desktop, tablet or mobile).

For each entry, basic information about the respective protein are provided, including the source type (Table 1), in addition to cross-references to major publicly available databases for diseases and drugs (DrugBank [23, 24], OMIM [25], Orphanet [26]), 3D structures (PDB [18]), protein families (Pfam [27], PROSITE [28]), genes (EMBL [29], HGNC [30]), pathways (KEGG [31], Reactome [32]), interactions (IntAct [33], BioGrid [34], STRING [35]), subcellular localization and tissue expression (COMPARTMENTS [36], Human Protein Atlas [37, 38]) and proteins (UniProt [20], RaftProt [39]). Moreover, a CytoscapeJS [40] viewer is integrated for the visualization of the interactions between peripheral membrane proteins and their interaction partners (when information is available in UniProt).

## 3. RESULTS

We have constructed PerMemDB, a relational protein database, which, in version 1.3 (March 2019), contains 231770 proteins originating from 1009 eukaryotes. The database can be either searched or browsed by organism. Each record contains sequence information and cross-references to many publicly available databases, with data spanning from protein family annotation to disease. Moreover, when available, information about the interaction partners of each peripheral protein in the database was retrieved from UniProt and is shown in an interaction network, that allows the quick identification of functional modules centering peripheral membrane proteins. There are 41828 entries isolated from UniProt (with subcellular location “Peripheral membrane protein”), 189925 were identified using MBPpred and 2325 were designated as indirectly interacting peripheral membrane proteins (TM interactors).

### 3.1. User interface and website features

The PerMemDB database has a user-friendly interface that offers convenient ways to gain access to its data. From the navigation bar at the top of every page, users can either perform searches or browse the database contents. Searching can be performed using various search terms (e.g. protein or gene name) and results may be refined by source type (UniProt, MBPpred or Transmembrane Interactor) or status (reviewed for UniProt/Swissprot entries or unreviewed for UniProt/TrEMBL entries). While browsing PerMemDB, a user can have access to all entries for a specific eukaryotic reference proteome (Figure 2). Results can be further filtered using the “Search” field at the top right of the page. When the green “Show selected Entries” button is pressed, the user is transferred to a new page with data regarding the proteome of the organism they have selected. Results, retrieved from either ‘browsing by organism’ or ‘searching’ the database, are displayed in tables. At the end of each row direct links to protein entry pages are given, which the user can follow by clicking on the respective buttons. Moreover, a BLAST [41] search tool is incorporated for running BLAST searches against the database proteins, using one or more FASTA formatted sequences as input.

**Figure 2.**
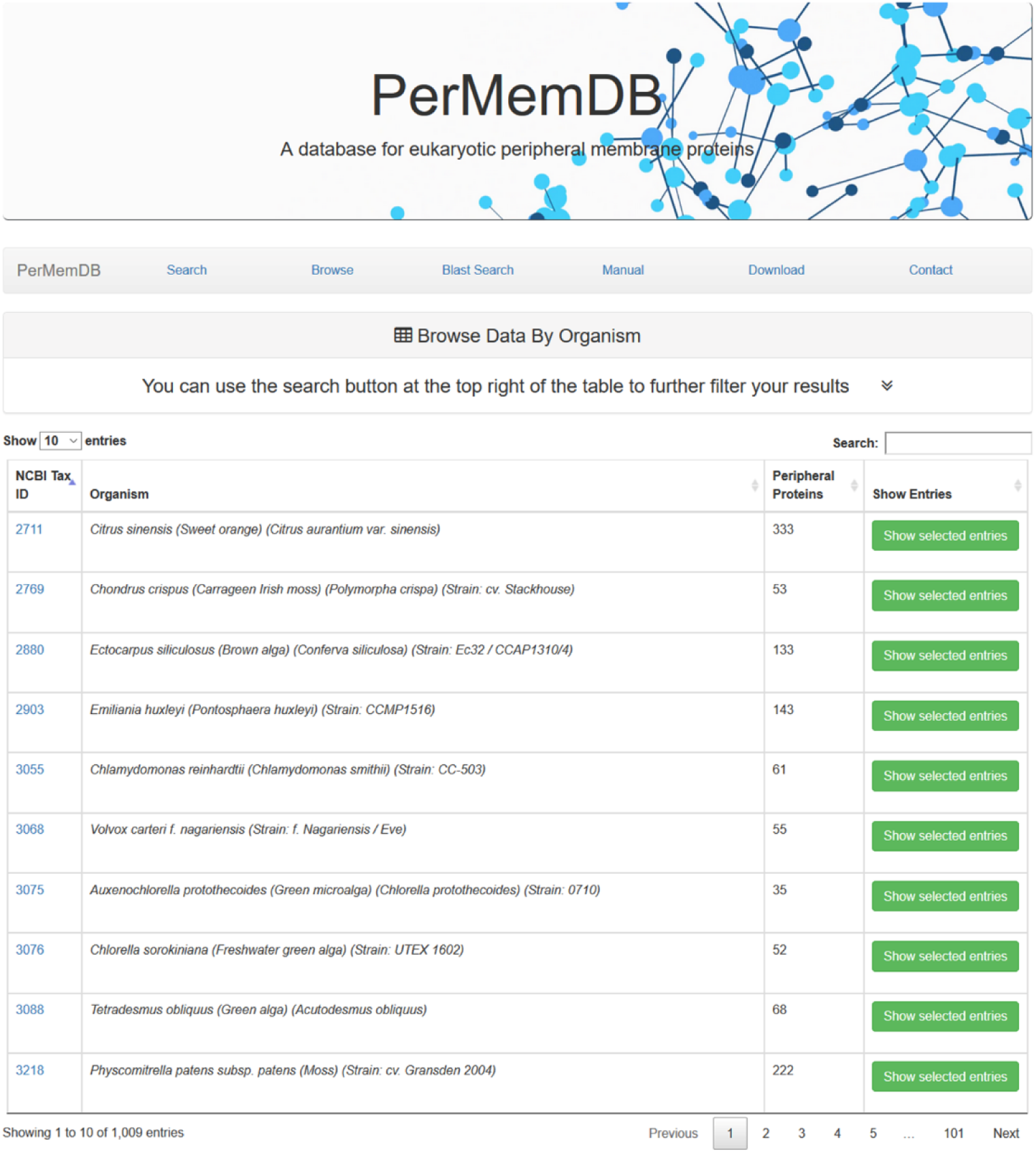
The ‘Browse’ Page of PerMemDB. Users can browse data stored in PerMemDB for a specific organism, whose proteome is listed as a eukaryotic reference proteome in UniProt. Non-specific searches can be performed using the ‘Search’ option at the top right corner of the data table. If a user presses the green ‘Show Selected Entries’ button all proteins from the selected organism are shown in a new page.

Apart from unique entries, the entire database is available for download by pressing the ‘Download’ button at the top navigation bar. PerMemDB is currently available in four formats (text, XML, JSON and FASTA). Finally, a ‘Home’ page for a short description and database statistics, a ‘Manual’ page explaining the functionalities of PerMemDB and a ‘Contact’ page with author contact information and a submission form to retrieve data from the scientific community, are also available.

Taking into consideration the biological and clinical significance of peripheral membrane proteins, our intention is to accurately represent all available information for this protein group through our repository. However, when the information source is all eukaryotic reference proteomes, this task can be challenging, and some data may be falsely filtered out during the necessary automated retrieval process. In an attempt to stay updated and be as comprehensive as possible, PerMemDB implements a user annotation feature. More specifically, in the contact page of our database, a form has been created dedicated explicitly to the submission of comments or data that has not been collected during the creation of the resource. Interaction with the users is of out-most importance to render this repository a useful tool for the scientific community. This process will allow us to incorporate valuable information regarding the sequence, the domains and the interaction networks of these proteins and thus, better annotate our entries and potentially improve our data retrieval protocol. It is our goal to implement all information gathered through this process in each database update, in addition to all data gathered using our automated pipeline.

### 3.2. Entry Pages

Database entries are generated dynamically via browsing, searching or through direct URL links. As shown in Figure 3, on the top of each page, data are available for download in four formats (text, XML, JSON and FASTA). Tables displaying basic information (e.g. protein name) and additional information (e.g. sequence) about each entry are shown on the left of each page. On the right, CytoscapeJS [40] is used to visualize the relationships between peripheral membrane proteins and their interaction partners. Links to RaftProt [39] are available for proteins with experimental evidence regarding their presence on lipid rafts, membrane substructures that compartmentalize cellular processes [42] and especially signal transduction processes. Moreover, direct links to the COMPARTMENTS database that contains information on protein subcellular localization from several sources (including manually curated literature, automatic text mining, and prediction methods) for seven model organisms, namely *Arabidopsis thaliana, Drosophila melanogaster, Homo sapiens, Mus musculus, Rattus norvegicus, Saccharomyces cerevisiae* and *Caenorhabditis elegans*, are given for entries associated with these organisms. On the bottom part of each page direct links to several external databases are also provided.

**Figure 3.**
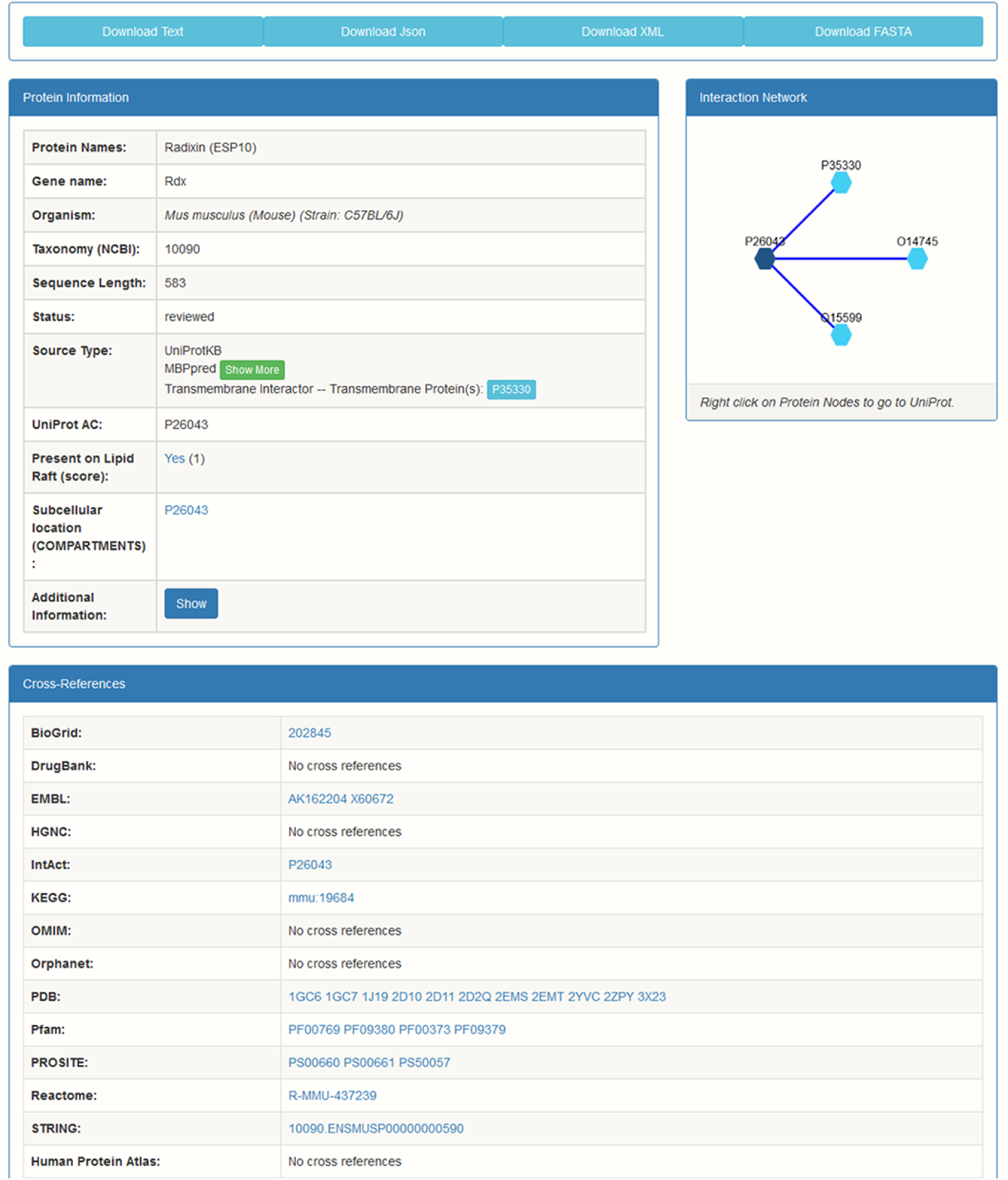
Detailed view of a protein entry of PerMemDB. The user can view basic information about the protein through the “Protein Information” panel. Additional information is provided by pressing the blue “Show” button. For entries retrieved through MBPpred information regarding the Membrane Binding Domains (MBDs) of the proteins are provided by pressing the green “Show More” button in the “Source Type” field. For peripheral proteins that have been identified as “non-transmembrane interactors of transmembrane proteins”, links to UniProt are provided for their transmembrane interaction partners (light blue button with UniProt AC). On the right, the relationships between peripheral membrane proteins and their interaction partners are shown when available. Upon clicking on the links at the bottom of the page users are transferred to the pages of the respective cross-references. All data can be downloaded in text, JSON, XML and FASTA format from the top of each entry page.

Information regarding the position and sequence of Membrane Binding Domains (MBDs) for proteins retrieved with the use of MBPpred is provided in the “Source Type” field of each protein entry. Moreover, in the same field, links to UniProt are given for transmembrane proteins designated as interaction partners of indirectly interacting peripheral membrane proteins (Figure 3).

### 3.3. Analysis of Database Data

#### 3.3.1 Quantification Analysis

With the aim of taking a closer look at the data stored in PerMemDB we performed a quantification analysis based on source type (Supplementary Table 1). At first glance, it is evident that the majority of data stored in the database were derived from the prediction method MBPpred (ca. 80%). Data originating solely from UniProt account for 18%, while a very small amount of data were designated as transmembrane interactors. Moreover, there is little cross-over between these three data sources. In particular, ∼1% of peripheral membrane proteins are found both in UniProt and by running MBPpred, and only 76 proteins were retrieved from all three sources. This was not unexpected though, since peripheral membrane proteins are generally understudied as a group, in comparison to other membrane protein groups. In addition, even though efforts for the annotation of this diverse group of proteins in human and specific model organisms have been carried out, things are extremely different for all other eukaryotes, which dominate the organisms populating our database (>95%) (Supplementary Table 1, Figure 4). Specifically, in the majority of eukaryotic reference proteomes in PerMemDB the number of predicted peripheral membrane proteins is more than tenfold that of the annotated proteins from UniProt. Moreover, databases dedicated to the recording of protein subcellular localization information are limited to only specific model organisms [36] or human [37], since data on the localization of proteins for other organisms are extremely scarce and difficult to detect. This underlines the fact that compared to generalized databases (like UniProt), PerMemDB presents a more complete coverage of available representatives for peripheral membrane proteins, since it provides information for a plethora of eukaryotic reference proteomes for which experimental, text-mined or prediction-based evidence is remarkably limited. Thus, our database could serve as a useful resource for the computational analysis and the clarification of the functional nature of this protein group.

**Figure 4.**
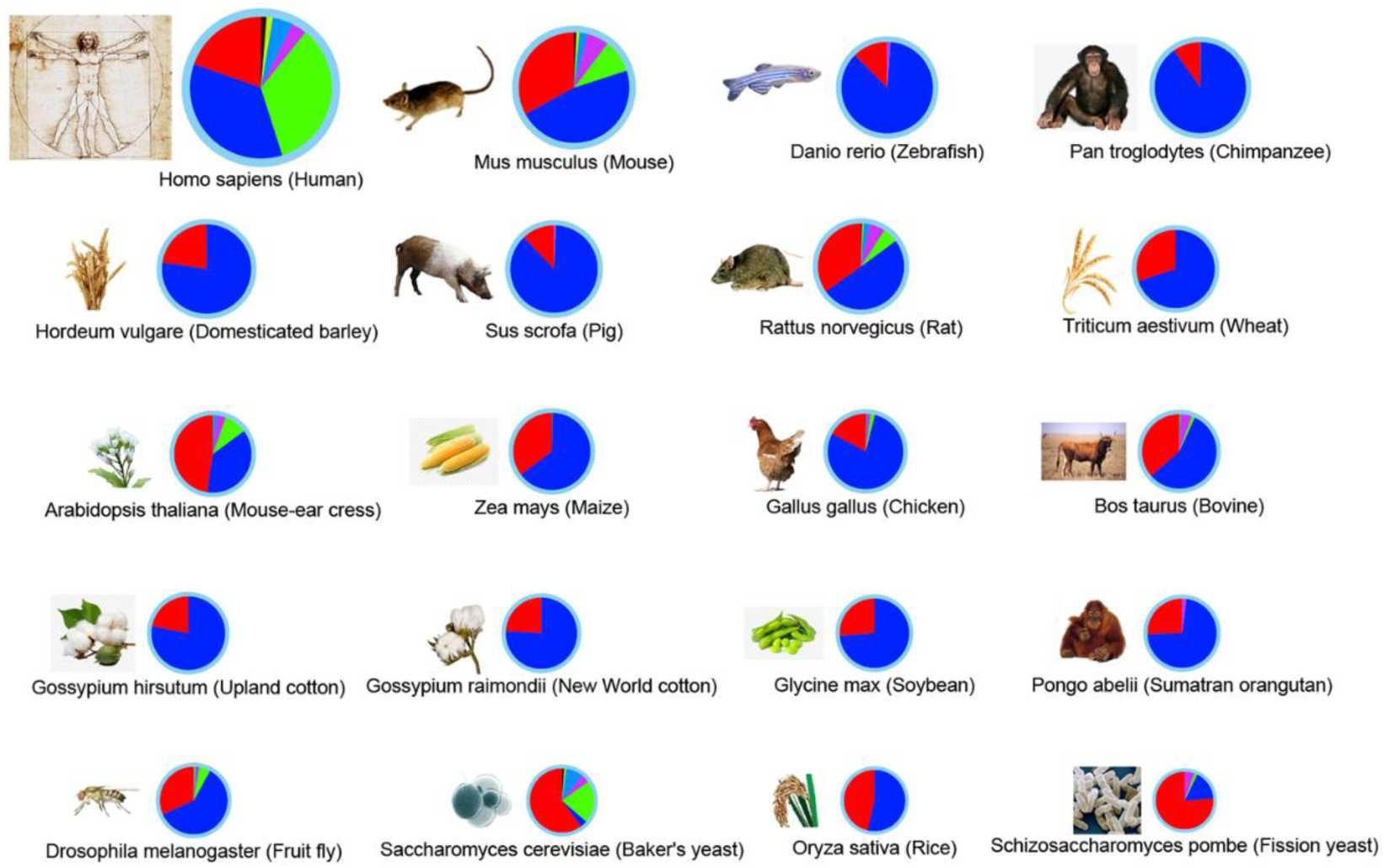
Twenty proteomes with the most representatives in UniProt. Pie charts are used to depict the number of peripheral membrane proteins for each proteome. The size of each pie chart is analogous to the total number of peripheral proteins detected in it. Data are color coded based on the subset in which the peripheral proteins belong to (See Table 1). Blue: UniProt_only, Red: MBPpred_only, Green: TMint_only, Magenta: UniProt_MBPpred, Light_blue: UniProt_TMint, Yellow: UniProt_MBPpred, Gray: UniProt_MBPpred_TMint. This image was created with the use of Cytoscape 3.7.0 [43].

In Figure 4 pie charts are used to depict the distribution of peripheral membrane proteins, based on data source, for the twenty proteomes with the most representative proteins in UniProt. The size of each pie chart corresponds to the total number of peripheral proteins, while the different slices represent the categories shown in Table 1. It is evident by the overrepresentation of blue and red colors in these charts that the data were retrieved mostly either with the use of MBPpred or from UniProt “Subcellular Location”. Only very well-annotated proteomes – human, mouse, rat, mouse-ear cress and baker’s yeast – show diversity regarding the source of peripheral membrane proteins.

A quantification analysis was also performed to examine the distribution of Membrane Binding Domains (MBDs) for proteins stored in PerMemDB that were retrieved via the application of MBPpred, since, as mentioned above, these entries comprise ∼80% of data in the repository. The two most abundant domains (that account for 45% of membrane binding domains in peripheral membrane proteins) are the Pleckstrin Homology (PH) domain and the C2 domain (Figure 5). Both domains are found in proteins that are either cytoskeleton constituents [44] or involved in cell signaling and enzymatic activities [45, 46], biological processes mainly performed by peripheral membrane proteins in cells [47]. Recently characterized MBDs, like KA1 and Golph 3, are detected only in a small number of peripheral membrane proteins (Figure 5).

**Figure 5.**
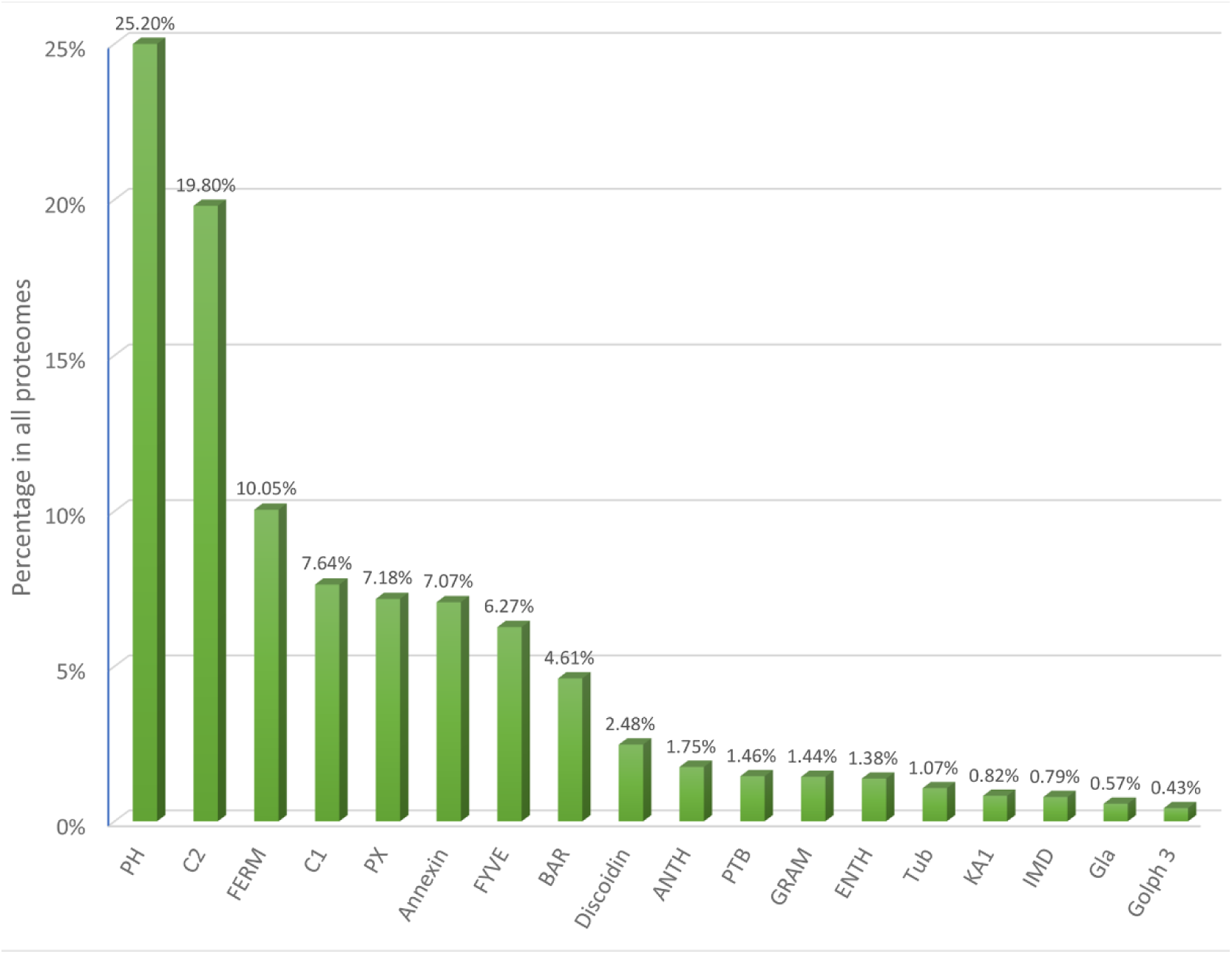
The distribution of Membrane Binding Domains (MBDs) in all protein entries retrieved via the application of MBPpred. The domains with the highest prevalence in peripheral membrane proteins are the well-studied PH and C2 domains, while recently identified domains like KA1 and Golph 3 are present in very small numbers.

#### 3.3.2 Functional enrichment analysis of ten selected proteomes

With the intention of investigating the functional roles of this diverse group of proteins, we analyzed the available data regarding the functions of these proteins for ten (10) selected proteomes (*Arabidopsis thaliana, Drosophila melanogaster, Danio rerio, Homo sapiens, Mus musculus, Rattus norvegicus, Saccharomyces cerevisiae, Gallus gallus, Bos taurus* and *Sus scrofa*). The functional enrichment tool incorporated in the Cytoscape stringApp [48] was used for this analysis. Detailed results are available in Supplementary Tables 2-11.

GO term enrichment analysis [49, 50] showed that proteins from these ten proteomes are located on membranes, vesicles or are involved in cytoskeleton organization, compartments where peripheral membrane proteins would be expected to be localized. Regarding their functions, they take part in catalytic activities and act mostly as kinases, which explains their tendency to participate in signal transduction processes. Moreover, considering their localization, it is only logical that these proteins participate in cell communication and vesicle-mediated transport. Thus, it is not surprising that peripheral membrane proteins are involved in many signaling pathways and endocytosis as indicated by the functional enrichment analysis of KEGG Pathways [31]. Finally, an enrichment analysis against data deposited in InterPro [51] and Pfam [27] revealed that these proteins contain, apart from known MBDs, other domains like PDZ [52], SH3, SH2 [53] and RhoGAP [54], which have all been observed to be present in peripheral membrane proteins, repeatedly.

#### 3.3.3 Analysis of pathogenic mutations on human peripheral membrane proteins

PerMemDB contains peripheral membrane protein data collected from different perspectives. Many inter- and intra-species applications that can contribute towards the better understanding of this group can be implemented. Considering the clinical significance of many peripheral membrane proteins [55, 56], we present here an application of PerMemDB for the study of the association between MBDs and disease variants in the subset of human proteins.

For this analysis, data on human genetic variations (“Polymorphisms” and “Disease” variants) were gathered from the UniProt database (release date: 13-02-2019). These variants were mapped on the sequences of human peripheral membrane proteins isolated from PerMemDB. A chi-square test was performed to get an estimate of differences in the emergence of “Polymorphisms” and “Disease” variants on regions that either do or do not contain MBDs.

The full dataset of missense Single Nucleotide Polymorphisms (SNPs) located on 341 out of the 3523 human peripheral membrane proteins consists of 2498 unique SNPs – 1471 “Disease” SNPs and 1027 “Polymorphisms”. From those, 434 SNPs are located on MBDs and 2064 on other regions. Considering the fact that these regions differ vastly in their length, all data had to be subjected to normalization, based on the length of each region.

Normalized data were subjected to chi-square testing to estimate the statistical difference between the frequency of “Disease” SNPs on MBDs and on other regions in peripheral membrane proteins. Results from this analysis show that there is a statistically significant difference (*p-value=0.029 < 0.05*) between the expected and observed “Disease” mutations in MBDs (Supplementary Table 12). These preliminary results indicate the importance of ‘structural’ protein information, like the topological profile of peripheral membrane proteins with MBDs, extracted from PerMemDB, in the evaluation of the implications of genetic variations in this protein group.

## 4. CONCLUSIONS

PerMemDB is currently the only repository that contains data dedicated exclusively to peripheral membrane proteins. The collection of data using three different methods allows a complete and extensive recording of all proteins that belong to this group. The existence of such a dataset can be very useful for large-scale proteomic analyses. As shown above, with the mutation analysis on human proteins, data deposited in PerMemDB can be valuable for disease association applications as well. Furthermore, taking into consideration the fact that PerMemDB contains data on a wide range of eukaryotes, it can serve as a valuable tool for evolutionary analyses, either for the entire group of peripheral membrane proteins or for membrane binding peripheral proteins with specific domains. The database is available for download for those who would like to access an updated and annotated dataset of peripheral membrane proteins for their research purposes, both in human readable text format and XML or JSON formats for programmatic access to the data.

Our goal is to keep the database up-to-date with biannual updates. Moreover, we aim to add novel membrane binding protein families to the MBPpred algorithm – if they are described in the scientific literature – which will allow for a more comprehensive resource, in the future. Finally, we hope that when more extensive studies on peripheral membrane proteins in prokaryotes become available, we will be able to populate the database with information about these organisms as well. To date, the role of prokaryotic proteins with domains that have membrane lipid-binding proteins in eukaryotes has not been revealed yet and in general, information on peripheral membrane proteins for these organisms is particularly limited. It is our hope that PerMemDB will aid the scientific community, towards gaining a profound understanding of this important group of proteins.

PerMemDB is available at http://bioinformatics.biol.uoa.gr/db=permemdb.

## ACKNOWLEDGEMENTS

The authors thank the National and Kapodistrian University of Athens for granting access to university premises and equipment.

## Author Contributions

Study design: KCN, GNT, VAI; Conceptualization: KCN, GNT; Database design and development: KCN; Data Curation: KCN, GNT; Web Application Design and Development: KCN; Web Application Quality Assurance: KCN, GNT, VAI; Writing - original draft: KCN; Writing - review and editing: KCN, GNT, VAI; Supervision: VAI.

## ABBREVIATIONS

MBD(s): Membrane Binding Domain(s)
MBP(s): Membrane Binding Protein(s)
pHMM(s): profile Hidden Markov Models
TM: transmembrane

